# Medication-Invariant Resting Aperiodic and Periodic Neural Activity in Parkinson’s Disease

**DOI:** 10.1101/2023.05.08.539920

**Authors:** Daniel J McKeown, Manon Jones, Camilla Pihl, Anna Finley, Nicholas Kelley, Oliver Baumann, Victor R. Schinazi, Ahmed A. Moustafa, James F. Cavanagh, Douglas J Angus

**Affiliations:** School of Psychology, Faculty of Society and Design, Bond University, Gold Coast, Queensland, Australia; Institute on Aging, University of Wisconsin-Madison, WI 53706, USA; School of Psychology, University of Southampton, United Kingdom; Department of Psychology, University of New Mexico, Albuquerque, NM 87106, United States

**Keywords:** Parkinson’s disease, EEG, cluster-based permutation tests, excitatory and inhibitory (E:I) balance, aperiodic activity

## Abstract

Parkinson’s Disease (PD) has been associated with greater total power in canonical frequency bands (i.e., alpha, beta) of the resting electroencephalogram (EEG). However, PD has also been associated with a reduction in the proportion of total power across all frequency bands. This discrepancy may be explained by aperiodic activity (exponent and offset) present across all frequency bands. Here, we examined differences in the eyes-open and eyes-closed resting EEG of PD participants (*N* = 26) on and off medication, and age-matched controls (CTL; *N* = 26). We extracted power from canonical frequency bands using traditional methods (total alpha and beta power) and extracted separate parameters for periodic (parameterized alpha and beta power) and aperiodic activity (exponent and offset). Cluster-based permutation tests over spatial and frequency dimensions indicated that total alpha and beta power, and aperiodic exponent and offset were greater in PD participants, independent of medication status. After removing the exponent and offset, greater alpha power in PD (vs. CTL) was only present in eyes-open recordings and no reliable differences in beta power were observed. Differences between PD and CTLs in the resting EEG are likely driven by aperiodic activity, suggestive of greater relative inhibitory neural activity and greater neuronal spiking. Our findings suggest that resting EEG activity in PD is characterized by medication-invariant differences in aperiodic activity which is independent of the increase in alpha power with EO. This highlights the importance of considering aperiodic activity contributions to the neural correlates of brain disorders.

## INTRODUCTION

More than 10 million people worldwide are currently diagnosed with Parkinson’s Disease (PD; Marras et al., 2018), experiencing significant motoric (e.g., tremors, impaired gait, rigidity, bradykinesia) and non-motoric (e.g., depression, loss of smell, emotional dysregulation, increased impulsivity) symptoms. Furthermore, the disease also affects carers, family members and friends due to the significant physical and emotional burden associated with caring for the individual with PD (Blaszczyk, 2016; Marras et al., 2018).

Coinciding with the gradual loss of dopaminergic neurons in the substantia nigra that contribute to motoric symptoms, PD is also associated with atypical neural oscillations and atypical levels of excitatory and inhibitory neurotransmitters (Little & Brown, 2014; O’Gorman Tuura et al., 2018; Weinberger et al., 2006). These alterations in dopaminergic function are thought to cause widespread pathological neural oscillations across canonical frequency bands (e.g., delta; 1 – 4 Hz, theta; 4 – 7 Hz, alpha; 7 – 13 Hz, beta; 13 – 30 Hz, gamma; >30 Hz) during tasks (George et al., 2013; Singh et al., 2018) and in resting state recordings (Olde Dubbelink et al., 2013). Individuals with PD show elevated total and relative beta compared to controls, with the most prominent differences typically observed over medio-central sites corresponding to the motor cortex. Although most studies have found that resting-state alpha is elevated in PD (Stoffers et al., 2007), others have reported a decrease (Caviness et al., 2016).

Furthermore, abnormal oscillations (e.g., beta power, alpha/theta ratios) have been linked to several factors related to the severity of PD (Miladinović et al., 2021). These include disease stage, disease progression, and the severity of cognitive impairment (Jaramillo-Jimenez et al., 2021; Olde Dubbelink et al., 2013). In addition, pharmacological treatments alter oscillatory activity in PD. Specifically, cortical and subcortical alpha and beta power decrease following the administration of Levodopa (a dopamine precursor), and are associated with a reduction in symptom severity (Cao et al., 2020; Kuhn et al., 2006; Kuhn et al., 2009).

Recent studies have highlighted that these differences in canonical frequency power could be underpinned (fully or partially) by aperiodic (i.e., non-oscillatory) signal activity. Aperiodic activity, also called 1/*f* activity since its power decreases as a function of frequency, was historically not considered to be physiologically meaningful or likely to impact estimates of oscillatory activity (Donoghue, Haller, et al., 2020). Recent evidence suggests otherwise, and highlights its potential importance for interpreting EEG correlates of brain disorders, including PD.

First, aperiodic activity is physiologically meaningful. The aperiodic exponent (i.e., the slope of the power spectra) has been shown to reflect the balance of excitatory and inhibitory (E:I balance) neural activity in both cross-sectional (Brady & Bardouille, 2022), animal model (Gao et al., 2017), and human experimental studies (Waschke et al., 2021). Here, smaller (i.e., flatter) exponents are observed following manipulations that increase excitatory activity and/or decrease inhibitory activity, and larger (i.e., steeper) exponents have been observed following manipulations that increase inhibitory neural activity and/or decrease excitatory activity. Whereas, the size of the aperiodic offset (i.e., the height of the power spectra) is directly associated with the rate of neural spiking (Manning et al., 2009), and with blood oxygen dependent levels during task performance (Winawer et al., 2013). Second, aperiodic activity varies systematically and semi-independently of oscillations. The aperiodic exponent has been found to vary as a function of age (Finley et al., 2022; Hill et al., 2022; Voytek et al., 2015), clinical status (Karalunas et al., 2022; Shuffrey et al., 2022), and cognitive function (Cross et al., 2022; Immink et al., 2021; Ostlund et al., 2021). Third, these systematic differences in aperiodic activity could act as a substantial confound and create the appearance of systematic differences in oscillations in total power and ratio-based measures (Donoghue, Dominguez, et al., 2020; Donoghue, Haller, et al., 2020). As such, traditional quantification methods, including those that rely on the calculation of ratios, can result in a misestimation and misattribution of aperiodic activity and periodic activity. Using EEG to determine the nature of aperiodic activity in PD offers unique insights into the pathophysiology of the disease and enhance the diagnostic potential of EEG.

The ability to infer E:I balance and neural spiking noninvasively is particularly important for understanding the neuropathology of PD. Recent work has shown that PD is also associated with alterations in GABAergic neurons and activity (Błaszczyk, 2016; O’Gorman Tuura et al., 2018) and that pharmaceutical treatment of these alterations (i.e., dopaminergic medications) may impact aperiodic activity by rebalancing E:I ratios (Belova et al., 2021; Wang et al., 2022). Convergent evidence from subdural electrophysiology (i.e., deep-brain stimulation, DBS; Belova et al., 2021; Darmani et al., 2023; Kim et al., 2022; Martin et al., 2018), neuroimaging, computational modelling (Moustafa et al., 2008), and drug trial (Song et al., 2021) studies suggest that alterations in E:I balance contribute to impairments and alterations in the behaviour of individuals with PD. Other studies have found that PD also alters the rate of neural spiking, particularly in the subthalamic nucleus (STN; Shimamoto et al., 2013), with the rate of spiking increasing with symptom severity and disease progression (Remple et al., 2011).

Considering this, the current study examined whether PD and sex- and age-matched healthy controls (CTLs) differ in resting aperiodic activity, and the extent to which the parametrization of oscillatory activity, can account for differences in canonical alpha and beta frequency bands between PD and CTLs. Using unpublished resting state data collected during previous studies (Cavanagh et al., 2018; Cavanagh et al., 2017), we examine variation in the EEG spectra between PD on and off medication and CTLs.

## METHODS

### Participants

Twenty-eight participants with PD were recruited from the Albuquerque, New Mexico community, and were paid $20 per hour for their participation. An equal number of CTL participants were taken from a larger sample reported elsewhere (Cavanagh et al., 2017). The University of New Mexico Office of the Institutional Review Board approved the study and all participants provided informed consent. Two PD and two CTL participants had insufficient clean EEG data and were excluded from analysis. EEG recordings were obtained from the PD participants on two occasions: once while on dopaminergic medication (ON), and once after a 15-hour overnight withdrawal from the dopaminergic medication (OFF). Demographic and diagnostic details for the retained participants (26 OFF/ON; 26 CTL) are presented in Table 1.

**Table 1.** Descriptive statistics for PD and CTL participants

|  | PD (n = 26) | CTL (n = 26) |
| --- | --- | --- |
| Age | 70.00 (8.26) | 69.50 (9.53) |
| Sex = Male (%) | 15 (57.7) | 16 (61.5) |
| Yrs Ed.Self | 17.42 (3.29) | 16.68 (3.04) |
| BDI | 7.08 (4.96) | 4.96 (4.82) |
| MMSE | 28.69 (1.05) | 28.85 (0.97) |
| NAART | 46.15 (8.79) | 46.88 (7.57) |
| Total Daily LED | 693.31 (446.20) | - |
| Yrs. Since Diagnosis | 5.46 (4.33) | - |
| UPDRS ON | 21.42 (10.18) | - |
| UPDRS OFF | 23.12 (8.62) | - |
| UPDRS DIFF | -1.69 (6.98) | - |

*LED, L-Dopa equivalence dose in mg; BDI, Beck Depression Inventory; MMSE, Mini Mental State Exam; NAART, North American Adult Reading Test; UPDRS, United Parkinson's Disease Rating Scale (motor); ON/OFF, when on or off medication respectively. Data is presented as means and SD.*

### Experimental procedure

The procedures used in the current study are described in detail elsewhere (Cavanagh et al., 2018). PD participants visited the lab on two occasions, one-week apart. At one visit, PD participants were on medication, while on the other visit they were off medication. The order of ON and OFF visits was balanced across PD participants, with 13 PD participants on medication during their first visit and 13 PD participants off medication during their first visit. Recordings for both PD and CTL participants were collected at 9AM. Neuropsychological assessments and questionnaires were completed by PD participants during their ON visit. In addition to the auditory oddball (Cavanagh et al., 2018) and value of volition (Cavanagh et al., 2017) tasks completed by PD and CTL participants, one-minute of eyes-closed (EC) and eyes-open (EO) resting state EEG was recorded during each lab visit. These recordings taken prior to the experimental tasks. For these recordings, participants were asked to sit and rest quietly, first with EC, and then again with EO. All participants completed the EC and EO recordings in the same order.

### Physiological recording and processing

Continuous EEG was recorded using 64 active Ag/AgCl electrodes, arranged according to the international 10/20 system. EEG signals were sampled at 500 Hz and online band-pass filtered between 0.1 and 100 Hz using a Brain Vision amplifier. CPz and Afz were used as the reference and ground respectively. Although vertical electrooculogram (VEOG) was recorded from auxiliary electrodes, these data were discarded during preprocessing. Preprocessing of EEG data was conducted using MATLAB (v2019a) and EEGLAB (v2021.0; Delorme & Makeig, 2004). Resting data were first referenced to the average of all scalp electrodes, and CPz was recreated. Data were then passed through the PREP pipeline, which we used to automatically identify and interpolate bad channels, reduce line-noise, and create a robust average reference (Bigdely-Shamlo et al., 2015). PREP processed data were high-pass (0.1 Hz) and notch filtered (60 Hz) to remove any remaining low-frequency and line-noise. We then used independent components analysis (ICA; Jung et al., 2000) and the Multiple Artifact Rejection Algorithm (MARA; Winkler et al., 2011; https://github.com/irenne/MARA) plugin to automatically identify and remove ocular, muscular, cardiovascular, and electrode artifacts. EC and EO data were then segmented into 2000ms segments overlapping by 50%. Segments in which the voltage of any scalp channel deviated by ±150 μV were marked as artifactual and excluded from analysis. Two participants with <50% of non-artifactual EC or EO data during any lab visit were also excluded from analysis.

Retained participants had an average of 97.05% (SD = 2.59%) non-artifactual data for EC recordings, and 97.43% (SD = 3.20%) for EO recordings. We then extracted the power spectra for EC and EO data from each channel for each participant using a Fast Fourier Transform (FTT) with a 2000ms Hamming window, overlapping by 50%.

### Parameterization of power spectra

We first extracted alpha and beta power using the traditional approach of taking the average power of the spectra between the canonical frequency bands, hereafter referred to as total alpha and total beta. To parameterize the power spectra into its aperiodic and periodic components, we used the algorithm implemented in the fitting oscillations & 1/*f* toolkit (FOOOF/specparam package; Donoghue, Haller, et al., 2020; https://github.com/fooof-tools/fooof). The FOOOF algorithm uses an iterative process to quantify both non-oscillatory aperiodic activity – which follows a 1/*f*-like power distribution – and oscillatory activity that overlaps with the underlying 1/*f-*like power distribution. The algorithm initially fits an aperiodic component to a power spectrum, which models both the exponent (i.e., the slope) and the offset (i.e., the height) of the spectra. To identify oscillations, the algorithm then regresses out the initial aperiodic component from the power spectra, and iteratively detects peaks using Gaussian modelling in the now flattened spectra, stopping this process when a predetermined number of peaks has been reached. After combining and refitting the oscillatory peaks, the algorithm then refits the aperiodic component, combines both oscillatory and aperiodic fits, and estimates the goodness of fit between the original power spectra and the algorithm derived model.

In the current study, and in keeping with previous research and recommendations (Finley et al., 2022; Ostlund et al., 2021), we parameterized the spectra from 2 – 40 Hz, using the following algorithm settings: peak width limits: 1-8; max number of peaks: 8; minimum peak height: 0.1; peak threshold: 2 SD; and aperiodic mode: fixed. For each channel by condition by participant combination, this approach yielded four parameters of interest: the aperiodic exponent, the aperiodic offset, parameterized alpha power, and parameterized beta power. Goodness-of-fit was evaluated using the mean *R*^*2*^ for each participant across EC and EO conditions. Fits were considered bad if they were < 3SD below the mean fit across all channels, conditions, and participants. One CLT participant in the EO condition had a mean fit below the threshold of *R*^*2*^ = 0.939. As the inclusion or exclusion of this participant did not substantially alter any results, we opted to retain them in the analysis^1^.

### Statistical analysis

Differences between ON, OFF, and CTL participants in aperiodic and periodic activity were examined using non-parametric cluster-based permutation testing as implemented in MNE. This approach adjusts for Type 1 error rate increases that are intrinsic to making within-or between-subjects comparisons across multiple electrodes.

Comparisons between ON and CTL and OFF and CTL were implemented using independent samples *t*-tests. Comparisons between ON and OFF were implemented using a 1-sample *t*-test performed on subject and electrode level difference scores between ON and OFF conditions. For each of the cluster-based tests, we used a cluster threshold of *p* < 0.05 with 10,000 Monte Carlo permutations. Consistent with previous studies, we report the maximum *t* statistic within each cluster, the Cohen’s *d* averaged across all electrodes within each cluster, and the *p* value for the cluster (Merkin et al., 2023). If analyses did not yield any significant clusters, we reported the maximum *t* statistic and Cohens *d* averaged across all electrodes and omit any cluster *p* value.

## RESULTS

### Aperiodic activity

First, we explored differences in the aperiodic exponent (Figure 1A-C) and aperiodic offset (Figure 1D-F). Cluster-based tests indicated that PD tended to be associated with significantly larger (i.e., steeper) exponents. We observed significant differences between OFF and CTLs in EC (*t*_max_ = 3.42, *d*_mean_ = 0.73, *p* = 0.024) but not in EO (*t*_max_ = 2.52, *d*_mean_ = 0.64, *p* = 0.060). Differences in the exponent between ON and CTLs were observed in EO (*t*_max_ = 2.82, *d*_mean_ = 0.66, *p* = 0.033) and EC (*t*_max_ = 3.13, *d*_mean_ = 0.70, *p* = 0.014). There were no significant clusters when comparing ON with OFF in either EO (*t*_max_ = 1.93, *d*_mean_ = 0.00) or EC (*t*_max_ = 1.99, *d*_mean_ = 0.04). Differences in the exponent between CTL and ON and CTL and OFF participants were primarily observed over medial central locations, with differences only extending to parietal locations for EC metrics.

**Figure 1.**
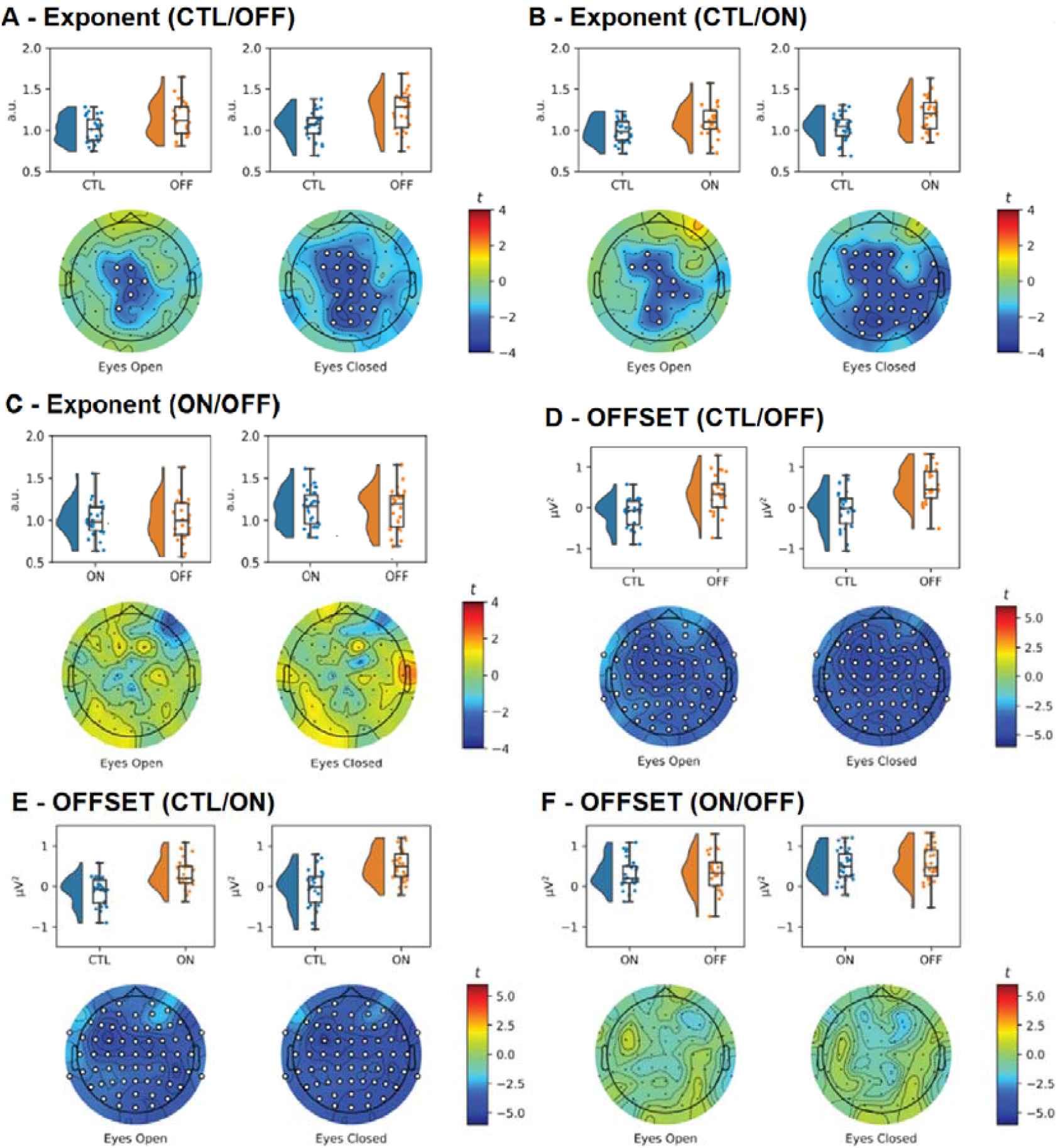
Raincloud and topographic plots comparing aperiodic activity in Parkinson’s disease (PD) and sex- and age-matched healthy controls (CTLs). Raincloud plots include the smoothed distributions, boxplots, and individual data points included in each of the analyses. Topographic plots depict the scalp distribution of the t-statistic for the tests comparing CTLs with off and on medication PD (OFF/ON), and ON with OFF. Electrodes with a t-statistic that exceeded the cutoff are masked in white. Panes A-C show differences in the aperiodic exponent for CTLs vs OFF (A), CTLs vs ON (B), and ON vs OFF (C), with greater (i.e., steeper) exponents in OFF/ON participants compared to CTLs. Panes D-F show differences in the aperiodic offset for CTLs vs OFF (D), CTLs vs ON (E), and ON vs OFF (F) participants, with greater offsets for OFF/ON participants compared to CTLs (n = 52).

There were clear, scalp-wide differences between PDs and CTLs in the aperiodic offset, with larger (i.e., higher) offsets observed for OFF vs. CTL (EO: *t*_max_ = 4.89, *d*_mean_ = 1.00, *p* < .001; EC: *t*_max_ = 5.01, *d*_mean_ = 1.12, *p* < .001), and ON vs. CTL (EO: *t*_max_ = 5.11, *d*_mean_ = 1.00, *p* < .001; EC: *t*_max_ = 5.05, *d*_mean_ = 1.10, *p* < .001). As with the aperiodic offset, there were no significant differences between ON and OFF (EO: *t*_max_ = 1.82, *d*_mean_ = 0.11; EC: *t*_max_ = 1.96, *d*_mean_ = 0.12).

### Alpha power

Next, we explored the differences in total alpha power (Figure 2 A-C) and parameterized alpha power (Figure 3 A-C). There were widely distributed differences between PD and CTLs for total alpha power, with greater total alpha power observed in PDs compared to CTLs regardless of medication status or EO/EC condition (OFF vs. CTL EO: *t*_max_ = 3.39, *d*_mean_ = 0.73, *p* < .001; OFF vs. CTL EC: *t*_max_ = 3.86, *d*_mean_ = 0.72, *p* = .004; ON vs. CTL EO: *t*_max_ = 3.18, *d*_mean_ = 0.67, *p* = .001; OFF vs. CTL EC: *t*_max_ = 3.61, *d*_mean_ = 0.71, *p* = .005). There were no significant differences between ON and OFF in EO (*t*_max_ = 2.62, *d*_mean_ = 0.50, *p* = .129) or EC (*t*_max_ = 1.89, *d*_mean_ = 0.01) conditions. Figure 3 A-C depicts the same analyses but examines parameterized alpha power – oscillatory power after having removed the aperiodic component.

**Figure 2.**
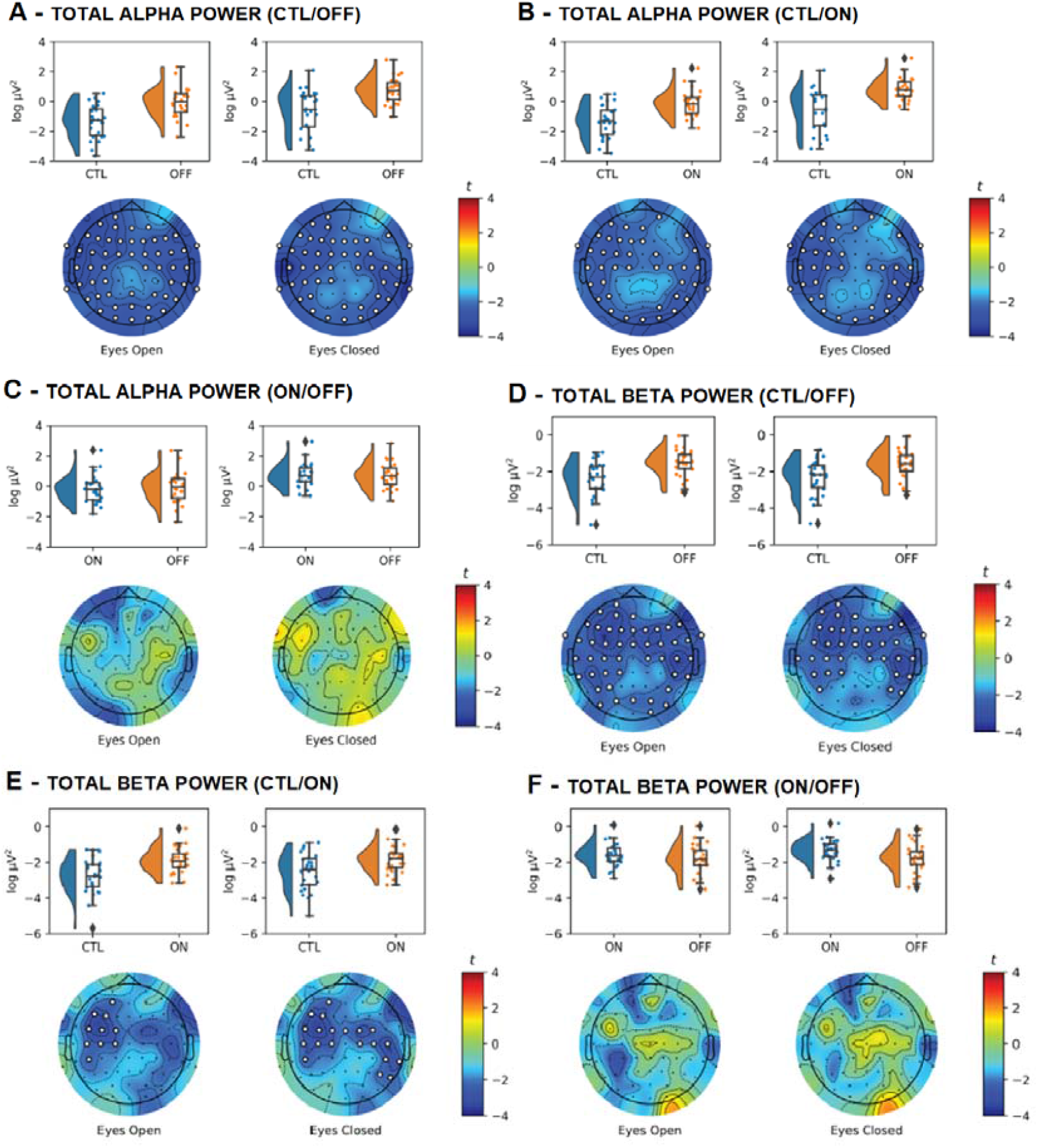
Raincloud and topographic plots comparing total alpha and total beta activity in Parkinson’s disease (PD) and sex- and age-matched healthy controls (CTLs). Raincloud plots include the smoothed distributions, boxplots, and individual data points included in each of the analyses. Topographic plots depict the scalp distribution of the t-statistic for the tests comparing CTLs with off and on medication PD (OFF/ON), and ON with OFF. Electrodes with a t-statistic that exceeded the cutoff are masked in white. Panes A-C show differences in total alpha power for CTLs vs OFF (A), CTLs vs ON (B), and ON vs OFF (C), with greater total alpha in OFF/ON participants compared to CTLs. Panes D-E show differences in total beta power for CTLs vs OFF (D), CTLs vs ON (E), and ON vs OFF (F) participants, with greater total beta power for OFF/ON participants compared to CTLs (n = 52).

**Figure 3.**
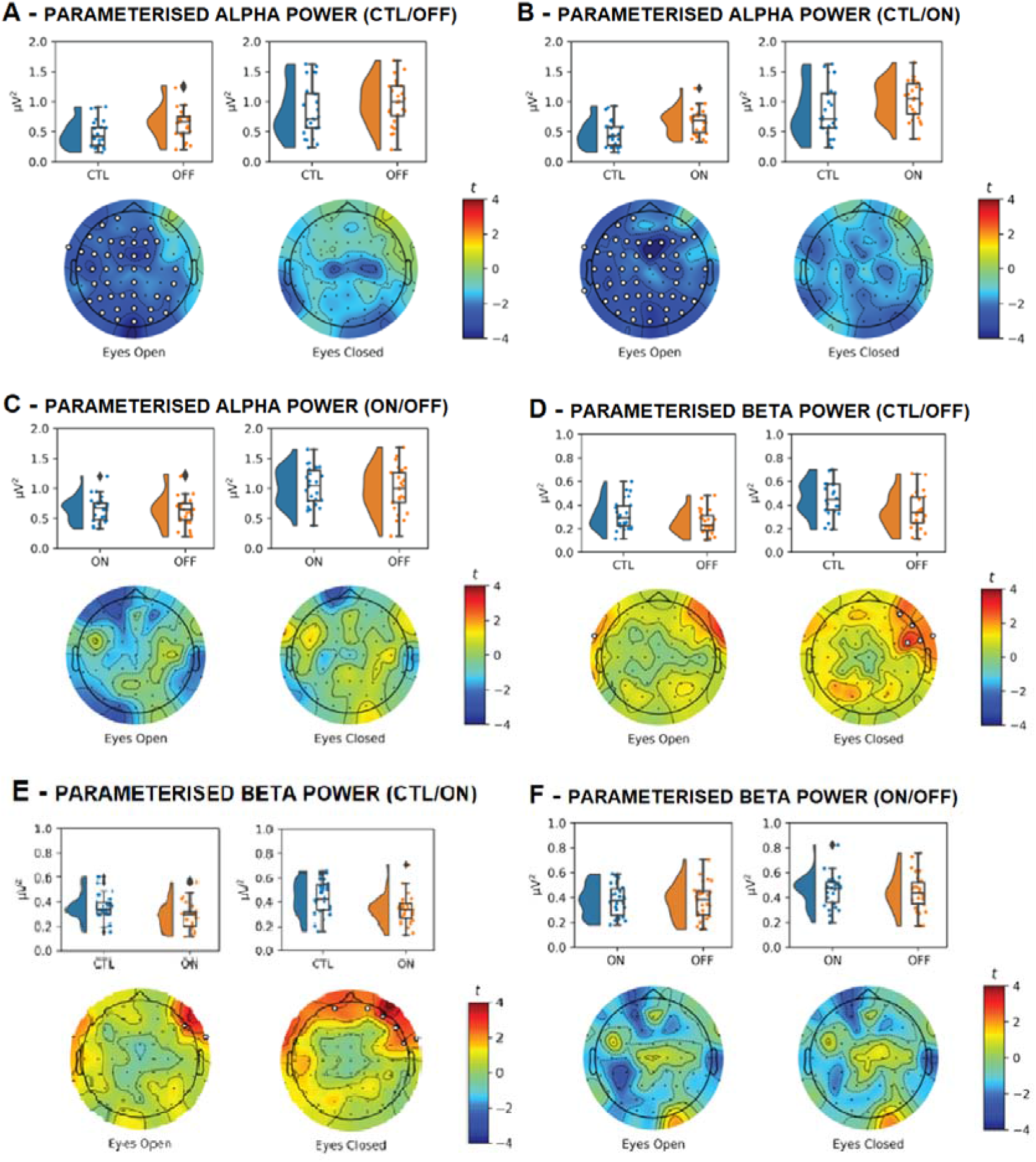
Raincloud and Topographic Plots Comparing Parameterized Alpha and Beta Activity in Parkinson’s disease (PD) and sex- and age-matched healthy controls (CTLs). Raincloud plots include the smoothed distributions, boxplots, and individual data points included in each of the analyses. Topographic plots depict the scalp distribution of the t-statistic for the tests comparing CTLs with OFF/ON, and ON with OFF. Electrodes with a t-statistic that exceeded the cutoff are masked in white. Panes A-C show differences in parameterized alpha power for CTLs vs OFF (A), CTLs vs ON (B), and ON vs OFF (C), with differences in power only observed between CTLs and OFF/ON in eyes-open conditions. Panes D-E show differences in parameterized beta power for CTLs vs OFF (D), CTLs vs ON (E), and ON vs OFF (F) participants (n = 52).

Although parameterized alpha power was greater for OFF and ON compared to CLTs, these differences were only significant in the EO condition (OFF vs CTL: *t*_max_ = 3.79, *d*_mean_ = 0.72, *p* = .012; ON vs CTL: *t*_max_ = 4.37, *d*_mean_ = 0.77, *p* = .006). There were no significant differences between OFF and CTL (*t*_max_ = 2.59, *d*_mean_ = 0.73, *p* = .287) and ON and CTL (*t*_max_ = 2.27, *d*_mean_ = 0.64, *p* = .134) in the EC condition, or between ON and OFF in EO (*t*_max_ = 1.55, *d*_mean_ = 0.00) or EC (*t*_max_ = 2.37, *d*_mean_ = 0.47, *p* = .695).

### Beta power

Lastly, we examined differences in total (Figure 2 D-F) and parametrized (Figure 3 D-F) beta power. Total beta power was consistently greater for PD participants vs CTLs. These differences were observed when comparing OFF with CTLs (EO: *t*_max_ = 3.47, *d*_mean_ = 0.72, *p* = .005; EC: *t*_max_ = 3.37, *d*_mean_ = 0.73, *p* = .009), and when comparing ON with CTLs (EO: *t*_max_ = 3.34, *d*_mean_ = 0.73, *p* = .031; EC: *t*_max_ = 3.39, *d*_mean_ = 0.71, *p* = .021). There were no differences in total beta as a function of medication status (EO: *t*_max_ = 2.55, *d*_mean_ = 0.47, *p* = .173; EC: *t*_max_ = 2.54, *d*_mean_ = 0.49, *p* = .347).

The cluster-based differences observed for total beta were not present when examining parametrized beta. Although there was greater parameterized beta for CTLs vs PD participants over right-fronto-polar electrodes in the EC conditions (Figure 3D and E), these clusters did not reach significance. That is, after removing the aperiodic component, there were no significant differences in beta power for CTL vs OFF in EO (*t*_max_ = 2.05, *d*_mean_ = 0.58, *p* = .453) or EC (*t*_max_ = 2.91, *d*_mean_ = 0.67, *p* = .077), CTLs vs ON in the EO (*t*_max_ = 2.42, *d*_mean_ = -0.64, *p* = .203) or EC (*t*_max_ = 2.92, *d*_mean_ = -0.64, *p* = .070). As with other measures, we did not observe a difference in parametrized beta or for ON vs OFF in the EO (*t*_max_ = 1.41, *d*_mean_ = 0.04) or EC (*t*_max_ = 1.82, *d*_mean_ = 0.08) conditions.

## DISCUSSION

In this study, we examined differences in alpha and beta band power before isolation from aperiodic activity (unparameterized) and when accounting for aperiodic activity (parameterized) in the resting state EEG of PD patients compared to CTL. We also explored the effect of medication status of PD patients on these EEG parameters. We found that individuals with PD showed higher exponents and larger offsets compared to CTL during both EO and EC conditions, regardless of medication status. Additionally, PD patients exhibited higher total alpha and beta power during both conditions, but these differences were reduced when accounting for aperiodic activity during both conditions for beta power, and only during EC for alpha power. These findings indicate that changes in canonical neural oscillatory activity in individuals with PD are primarily driven by changes in E:I balance favouring greater inhibition (aperiodic exponent) and greater neural spiking rate (aperiodic offset) when at rest.

### Aperiodic activity in Parkinson’s Disease

Converging evidence indicates that impairment in neurotransmitter (dopamine and GABA) balance contributes to alterations in the behaviour of individuals with PD (Song et al., 2021). Indeed, impairment in dopaminergic and GABAergic neuronal activity are prominent in PD (Blaszczyk, 2016; Meder et al., 2019; O’Gorman Tuura et al., 2018). Here, a reduction in dopamine concentrations in the substantia nigra pars compacta and subthalamic nucleus (STN), key components of the basal ganglia and integral for motor control, has been found to result in over-inhibition and slowing of movement (i.e., bradykinesia; Chu et al., 2017; Meder et al., 2019). Similarly, reductions in GABA concentrations in the basal ganglia, motor cortex, and occipital cortex are associated with motor (resting tremor, reemergent tremor, and bradykinesia; van Nuland et al., 2020) and non-motor (hallucinations; Firbank et al., 2018) symptoms, the severity of which is determined by the progression of PD. PD also alters the rate of neural spiking, particularly in the STN (Shimamoto et al., 2013), with the rate of spiking increasing with symptom severity and disease progression (Remple et al., 2011). Concerning GABAergic neuronal activity, these findings are not surprising given the major GABAergic pathways in the basal ganglia, in contrast to other subcortical structures (Caligiore et al., 2016). Despite discrepancies in GABA changes in PD being prominent, they are likely related to differences in disease status, the subtype of PD group under investigation (i.e., tremor-dominant vs. akinesia-dominant), PD symptoms under investigation (axial vs. cardinal), as well as medication status.

The quantification of non-oscillatory aperiodic activity provides insight into local and widespread excitatory and inhibitory neural activity (Finley et al., 2022). Furthermore, the ability to infer E:I balance (aperiodic exponent), rate of neural spiking (aperiodic offset), and underlying changes in neurotransmitter availability, is particularly important for understanding the neuropathology of PD (Gao et al., 2017). Of the few studies that have characterized changes in aperiodic activity in PD, investigation have primarily been localised to deep brain regions using DBS (i.e., STN; Belova et al., 2021; Darmani et al., 2023; Kim et al., 2022; Martin et al., 2018). In these studies, the aperiodic exponent positively correlated with DBS treatment at rest (greater exponent slope from 2 to 6 months of DBS; Darmani et al., 2023), negatively correlated with voluntary muscle contractions of the finger flexors and tibialis anterior muscle independent of medication status (reduced exponent slope indicating greater excitation; Belova et al., 2021), and correlated with dopamine depletion (Kim et al., 2022). Together, these findings highlight the physiological relevance of aperiodic activity in the activation of the STN in PD. Consequently, increased inhibition of the STN (indicated by a greater exponent slope) has been identified as a potential mechanism for the efficacy of long-term DBS in the treatment of PD and as a biomarker for disease progression in PD (Darmani et al., 2023; Kim et al., 2022; Martin et al., 2018).

Although changes in aperiodic activity during DBS provide evidence for the STN being a potential neural generator for aperiodic activity, its pertinence is limited by the surgical invasiveness of the procedure. As the only other current study assessing aperiodic activity noninvasively with scalp EEG in PD, Wang et al. (2022) determined differences in the aperiodic exponent and offset in 15 PD on and off medication, compared to 16 CTL. Although statistically underpowered, dopaminergic medication increased the exponent and offset in patients with PD compared to off medication, but they were not different from CTL. Furthermore, these differences were greatest in the bilateral central brain regions (C4, C3, CP5, and FC5 regions of the scalp EEG) and was concluded to be due to the rebalancing effect of dopaminergic medication on E:I balance in the basal ganglia. In the current study, a greater aperiodic offset was observed in PD individuals compared to CTL during EO and EC conditions, while the aperiodic exponent was only significant during the EC condition. Furthermore, these findings were sufficiently statistically powered and observed in a larger PD population than previously investigated (Wang et al., 2022), independent of medication status (Darmani et al., 2023), found throughout multiple brain regions, including the bilateral central regions previously documented (Wang et al., 2022), and aligned with those documented during DBS (Belova et al., 2021; Darmani et al., 2023). This provides evidence of the potential efficacy of using non-invasive scalp EEG as an alternative to invasive subdural electrodes and sensing-enabled implantable pulse generators when identifying disease and treatment biomarkers of PD.

### Periodic alpha and beta activity in Parkinson’s Disease

Periodic activity has traditionally been assessed by the quantification of neural oscillatory activity in canonical frequency bands (i.e., alpha and beta band activity) while not controlling for aperiodic activity. This is problematic, as underlying aperiodic activity may result in misinterpretation of periodic activity. Considering this, previous conclusions on the changes in alpha and beta activity in PD may in fact be due to changes in the exponent and/or offset of aperiodic activity. This likely contributes to the conflicting findings of periodic activity assessment in PD. Indeed, increases in the power and synchronisation of oscillations in the beta band are frequently reported (Canessa et al., 2016; Little & Brown, 2014; O’Gorman Tuura et al., 2018; Weinberger et al., 2006) but have also been reported to decrease in line with disease severity (Vinding et al., 2020). Whereas, alpha band power is low in bilateral parietooccipital locations in PD individuals with dementia (Yilmaz et al., 2020) and higher without dementia (Stoffers et al., 2007). This highlights the necessity for parameterisation of these alpha and beta bands to accurately reflect changes in oscillatory activity in PD.

Once alpha and beta activity are parameterised, mixed responses are documented in PD. Although total power and paramaterised power have been reported to be comparable in both alpha and beta bands (Belova et al., 2021; Wang et al., 2022), parameterised beta power has been reported to be more sensitive to changes in local field potential features (Belova et al., 2021) and is significantly more accurate at estimating the beta biomarker of PD symptom severity (Martin et al., 2018). Indeed, the correlation between aperiodic activity and motor symptoms of PD (bradykinesia and rigidity) is most evident once beta band power is isolated from the aperiodic exponent and offset (Martin et al., 2018). In the current study, total alpha and beta power were significantly greater in PD during both conditions and independent of medication status. Once isolated from aperiodic activity, parameterised alpha and beta power ceased to show significance. This indicates that the change in oscillatory activity between PD and CTL can be accounted for by a change in underlying aperiodic activity and not alpha and beta power. That is, individuals with PD exhibited a greater proportion of inhibitory neural balance and an increase in the rate of neural spiking in the current study, mechanisms of which are independent of periodic activity (Gao et al., 2017).

### Dopaminergic medication and EEG activity

In the current study, no differences in periodic nor aperiodic activity were observed in PD patients between on or off medication. This suggests that dopaminergic medication did not change EEG signatures in our PD cohort. Conversely, Wang et al. (2022) found that dopaminergic medication increased the aperiodic exponent and offset, specifically in the bilateral central brain regions, and that this aperiodic activity in ON did not differ with CTL. The increase in aperiodic activity was suggested to be due to the recruitment of inhibitory inputs from the globus pallidus external to the STN. To date, six main types of medications are available to alleviate symptoms of PD, including levodopa, dopamine agonists, inhibitors of enzymes that inactive dopamine (i.e., monoamine oxidase type B inhibitors [MAO B] and catechol-O-methyl transferase inhibitors [COMT]), anticholinergics, and amantadine (Connolly & Lang, 2014). Considering our PD population were also on dopaminergic medication, it is likely the variability in symptomatology whilst off medication contributed to the similarities in periodic and aperiodic activity between PD and healthy CTL in the current study. Alternatively, the presence of medication invariant alterations in EEG signatures of PD patients found in the current study provides evidence that EEG signatures may further serve to predict aspects of PD disease progression that are not detected in most common phenotypic assessments.

### Implications

Although our findings are consistent with those of DBS (Belova et al., 2021; Darmani et al., 2023; Kim et al., 2022; Martin et al., 2018), oscillatory measurements derived from EEG are hard to localize. While our study reveals atypical exponents in PD, we cannot say which structures acted as neural generators for aperiodic activity in our cohort. Nonetheless, the use of EEG provides a non-invasive approach to characterise aperiodic and periodic activity biomarkers in PD. This study provides evidence that the contribution of aperiodic activity to canonical alpha and beta band power needs to be considered carefully, and previous models using MEG/EEG need to be reevaluated. Indeed, differences in alpha and beta band power may be artifactual and instead driven by differences in the aperiodic exponent and offset.

## Conclusion

When parametrising alpha and beta band power derived from EEG at rest in individuals with PD and CTL, the increase in total alpha and beta power is in fact due to an increase in the aperiodic exponent (greater inhibition) and an increase in the aperiodic offset (greater neural spiking rate) and not in fact due to changes in paramaterised alpha or beta band power. Furthermore, changes in aperiodic activity were independent of medication status. These findings correlate with those of previous DBS findings, and in doing so, provide a non-invasive approach to assess underlying changes in E:I balance and PD biomarkers.

## DATA AND CODE AVAILABILITY STATEMENT

The raw data used in this study is available at http://predict.cs.unm.edu/downloads.php, Accession # d002. Processed data and for all analyses and plots are publicly available at https://github.com/MindSpaceLab/Parkinsons_Aperiodic_Periodic.

## AUTHOR CONTRIBUTIONS

Conceptualization: M.J., A.J.F., N.K., A.A.M., J.F.C., and D.J.A. Data curation: D.J.M., M.J., J.F.C., and D.J.A. Formal analysis: D.J.M., M.J., and D.J.A. Project administration: J.F.C. and D.J.A. Visualization: D.J.A. Writing - original draft: D.J.M., M.J., and D.J.A. Writing - review & editing: D.J.M., M.J., C.P., A.J.F., N.K., O.B., V.S., A.A.M., J.F.C., and D.J.A.

## ACKNOWLEDGEMENTS

We would like to express our gratitude to all the participants involved in our study, for their support and patience, and for contributing their time.

## Footnotes

1 Exclusion of this participant only changed one result: Once excluded, parametrized beta power in the EC condition over a fronto-polar cluster was significantly greater in CTL participants compared to ON participants (*t*_max_ = 3.13, *d*_mean_ = 0.71, *p* = 0.041). No other results were altered.

